# Gene Expression Meta-Analysis Reveals Aging and Cellular Senescence Signatures in Scleroderma-associated Interstitial Lung Disease

**DOI:** 10.1101/2023.11.06.565810

**Authors:** Monica M. Yang, Seoyeon Lee, Jessica Neely, Monique Hinchcliff, Paul J. Wolters, Marina Sirota

## Abstract

Aging and cellular senescence are increasingly recognized as key contributors to pulmonary fibrosis. However, our understanding in the context of scleroderma associated interstitial lung disease (SSc-ILD) is limited. To investigate, we leveraged previously established lung aging and cell-specific senescence signatures to determine their presence and potential relevance to SSc-ILD. We performed a gene expression meta-analysis of lung tissue from 38 SSc-ILD and 18 healthy controls and found markers (GDF15, COMP, CDKN2A) and pathways (p53) of senescence were significantly increased in SSc-ILD. When probing the established aging and cellular senescence signatures, we found epithelial and fibroblast senescence signatures had a 3.6-fold and 3.7-fold enrichment respectively in the lung tissue of SSc-ILD and that lung aging genes (*CDKN2A, FRZB, PDE1A, NAPI12)* were increased in SSc-ILD. These signatures were also enriched in SSc skin and associated with degree of skin involvement (limited vs. diffuse cutaneous). To further support these findings, we examined telomere length (TL), a surrogate for aging, in lung tissue and found independent of age, SSc-ILD had significantly shorter telomeres than controls in type II alveolar cells in the lung. TL in SSc-ILD was comparable to idiopathic pulmonary fibrosis, a disease of known aberrant aging. Taken together, this study provides novel insight into the possible mechanistic effects of accelerated aging and aberrant cellular senescence in SSc-ILD pathogenesis.

## 1. Introduction

Systemic Sclerosis (SSc) is an autoimmune disease characterized by diffuse fibrosis and vasculopathy that has the potential to affect nearly every organ system. Interstitial lung disease (ILD) is present in up to 90% of SSc patients with many presenting early in their disease course with variable trajectories.(1–3) Despite advances in the field, ILD remains the leading cause of SSc morbidity and mortality accounting for up to 35% of SSc related deaths and a 5-year mortality rate ranging between 18-44%.(4,5) Management of SSc-ILD remains challenging due to significant disease heterogeneity and lack of precision medicine approaches. An improved understanding of the molecular mechanisms underpinning SSc-ILD is needed to advance treatment in the field.

There is a growing interest in the role of aging and cellular senescence in the development and progression of pathologic lung remodeling pulmonary fibrosis. Multiple epidemiologic studies have shown age as a risk factor for lung fibrosis and portends a worse prognosis in lung disease.(6,7) Aged lungs have been shown to have shortened telomeres and increased expression of the cellular senescence markers p16 and p21.(8,9) Idiopathic pulmonary fibrosis (IPF), the most common form of ILD, is characterized by telomere dysfunction, senescence programming and is a disease of older age.(10–12) A defining feature in IPF is that alveolar type II cells exhibit telomere shortening, adopt features of senescent cells and lose their ability to proliferate.(13,14) These processes are increasingly implicated across several forms of ILD, including connective tissue disease ILDs.(15,16) Better understanding of these shared mechanisms and how they may relate to SSc-ILD, has prognostic and treatment implications for an otherwise rare and understudied disease.

Transcriptomic studies have examined gene expression changes during lung aging and senescence, establishing signatures that can be used to probe existing datasets. Utilizing bulk RNAseq of healthy lungs across the lifespan, Lee et al. showed both cellular senescence and profibrotic pathways are increased with age and identified a lung-specific aging signature.(8) The 22 gene signature was validated in the Gtex lung dataset with a subset of genes also found to be present in aging skin. DePianto et al. generated *in vitro* senescent cells to define an epithelial senescence, fibroblast senescence and consensus senescence signature which was recapitulated in a scRNAseq data from IPF and to a lesser degree SSc-ILD.(17) While these mechanisms have been examined in murine models and in other human fibrotic lung diseases, their relevance to SSc-ILD has yet to be established.

Studies of lung tissue in SSc-ILD are limited due to disease rarity and tissue scarcity. Meta-analysis of publicly available data provides a practical approach for understanding gene expression changes in SSc-ILD as well as performing cross tissue comparisons, such as skin and blood. To date, gene expression meta-analyses in SSc-ILD have taken a largely exploratory approach, primarily focused on gene expression changes at large and shared features between tissues.(18,19) However, by taking a hypothesis driven approach by investigating gene signatures relevant to pulmonary fibrosis in both *in vitro* and *in vivo* studies, we can obtain additional insight into the pathogenesis of SSc-ILD as well as other SSc organ involvement.

In this study, we performed a gene expression meta-analysis of lung tissue from SSc-ILD to examine if aging and senescence related mechanisms are present in the SSc-ILD compared to healthy controls. We leveraged established lung-specific aging and cellular senescence signatures that have yet to be examined in the existing SSc-ILD data. To validate our findings, we measured telomere length in SSc-ILD, IPF and healthy control lungs. Finally, given SSc is a disease associated with skin fibrosis, we also examined if these signatures may be relevant to SSc skin disease.

## 2. Materials and Methods

### 2.1. Gene expression meta-analysis pipeline

A schematic overview of the meta-analysis pipeline is presented in Figure 1A. Publicly available microarray data sets from Gene Expression Omnibus (GEO) were searched for keywords “scleroderma”, “systemic sclerosis” and “interstitial lung disease”. Samples were curated to only include lung tissue from adult patients meeting ACR 2019 criteria for SSc who carried a diagnosis of SSc-ILD. When a study included multiple tissue samples for a single subject, expression data from the lower lobe was favored given the predilection for disease in that area, and other samples were excluded to avoid overrepresentation of any individuals. Healthy controls from each respective study were also included. Clinical covariates including age, sex, and disease duration were not available for all individual samples and therefore not included in the study.

**Figure 1:**
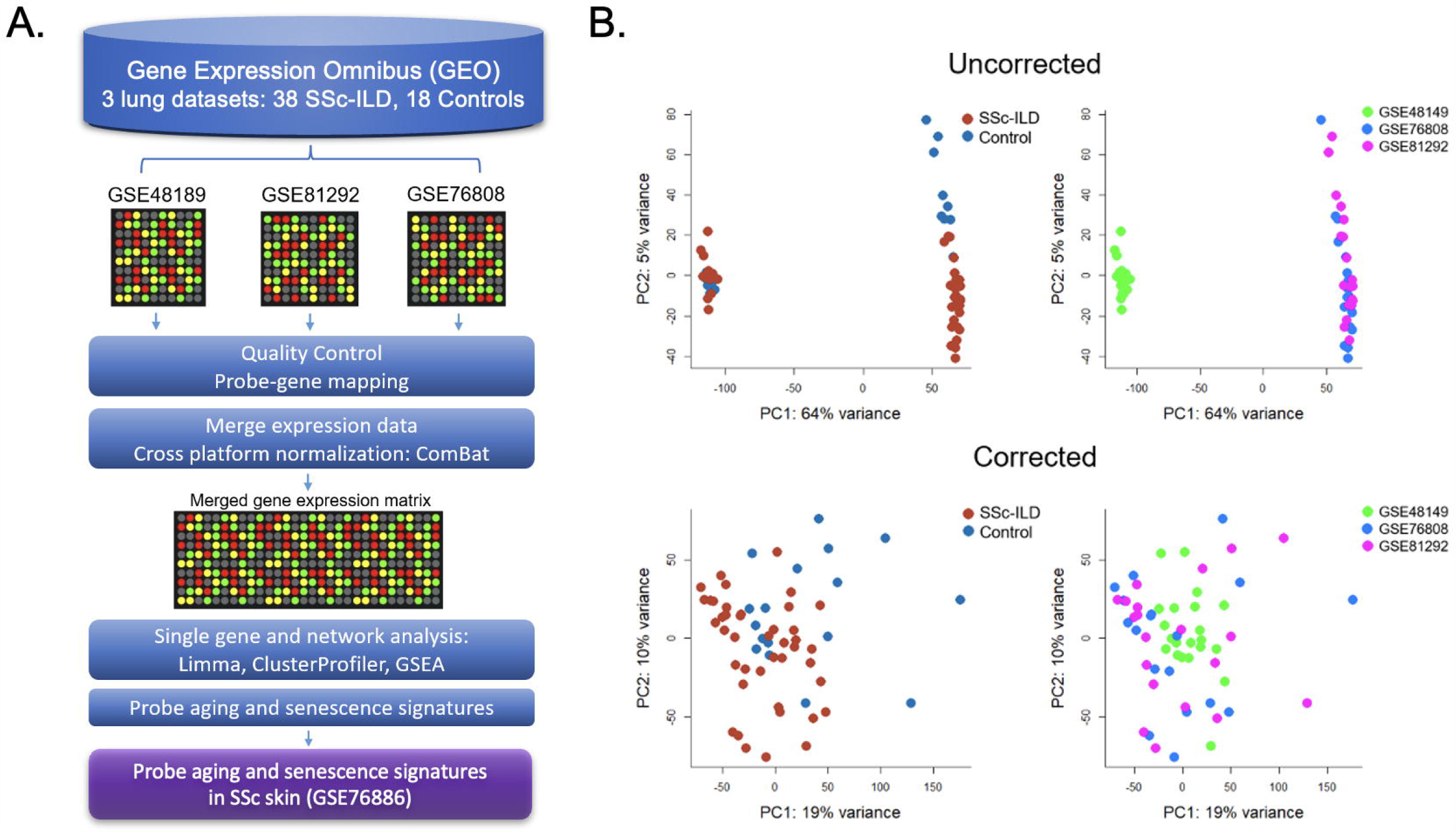
Gene expression meta-analysis pipeline (A) with PCA plots demonstrating successful merge after batch correction (B).

Raw data from each microarray data set was downloaded from GEO. Processing first took place for each individual dataset according to their unique microarray platform which included background correction, log2 transformation, normalization, and probe to gene mapping using R. LIMMA was utilized to process Illumina arrays, while SCAN.UPC from Bioconductor was utilized for Affymetrix arrays.(20,21) To be merged, expression data was mean-centered and reduced to genes common to all datasets. Cross study normalization and data set merging to create a single gene expression matrix was performed using ComBat, an empirical Bayes method.(22) Principal component analysis (PCA) and boxplots were used to evaluate the successful batch correction and to detect outliers.

### 2.2 Differential gene expression

Differential gene expression analysis was performed using LIMMA.(21) Genes considered differentially expressed (DEGs) between cases and controls were defined using a cutoff false discovery rate (FDR) *q*-value < 0.05 and fold change (FC) ≥ 1.2 or more. Unsupervised hierarchical clustering using the Ward algorithm was performed to generate heatmaps and identify gene expression clusters. Network analyses using Gene Ontology Enrichment Analysis and Gene Set Enrichment Analysis (GSEA) were conducted to identify genes and pathways significantly associated with SSc-ILD (p ≤0.05).(23,24)

### 2.3 Cell Type Enrichment

Cell type enrichment analysis was performed using xCell v1.1, a gene-signature based method, on the merged normalized data.(25) The existing compendium of cell type signatures were filtered to remove cell types not relevant to lung biology (i.e., osteoblasts, hepatocytes, keratinocytes, neurons) leaving 45 cell types of the immune, stromal, and epithelial compartments. The false discovery rate was controlled by the Benjamini-Hochberg method and the Wilcoxon test was used to determine differential cell enrichment between cases and controls using a threshold of adjusted *p* < 0.1. Significantly differentially enriched cell types were visualized with hierarchical clustering using the Ward algorithm.

### 2.4 Probing Aging and Senescence Signatures

Aging and senescence gene expression signatures have been established for the lung. Using bulk RNA-seq of in vitro senescent cells, DePianto et al. defined an epithelial senescence (227 genes), fibroblast senescence (117 genes), and consensus senescence signature (11 genes) (17). Utilizing bulk RNAseq of healthy lungs across the lifespan, Lee et al. identified a lung-specific aging signature including 16 upregulated genes that were validated in publicly available data of healthy controls (8). These signatures were used to determine whether aging and cellular senescence were enriched in SSc-ILD. For senescence, each cell-type specific signature was probed for overlapping genes within the merged gene expression matrix and visualized with unsupervised hierarchical clustering. Ratio of enrichment of each signature was determined using hypergeometric testing. For aging, overlapping genes were examined individually and gene expression between SSc and controls were compared using unpaired, two-sided t-test.

### 2.5 Telomere length measurement using TeloFISH

To further validate the findings of enhanced aging and cellular senescence in SSc-ILD, cell specific telomere length was measured utilizing telomere quantitative fluorescence in situ hybridization assay (TeloFISH) in lung tissue.(26) Lung tissue from 15 age and sex matched subjects including 5 SSc-ILD, 5 healthy controls, and 5 IPF, were fixed with 10% formalin overnight and embedded in paraffin and then sectioned to 4 um thickness. Tissue slides were deparaffinized in xylene and rehydrated in ethanol gradient series. After heat induced antigen retrieval, tissues were treated with 10mg/ml RNase solution and incubated with 0.3 μg/ml peptide nucleic acid (PNA) TelC-Cy3 probe (Panagene, Korea) in 70% formamide buffer, heated to 78°C for 10 min then overnight at 20°C. Following washing and blocking, tissues were incubated with primary rabbit-anti SPC antibody, followed by secondary antibody. Tissues were mounted with DAPI. Images were acquired using a Zeiss Axio Imager 2 microscope and telomere signal intensity was quantified in a blinded manner using MetaMorph imaging analysis software (Molecular Devices, Sunnyvale, CA). Pairwise comparisons of telomere length were made between groups using Kruskall Wallis test.

### 2.6 Aging and Senescence signatures in SSc skin

To examine if aging signatures were also relevant in the skin in SSc, we used a previously published gene expression dataset from the skin (GSE76886). Sample processing and sequencing has been described in detail previously.(27,28) PCA was used to visualize outliers. Metadata including SSc disease subtype (limited cutaneous vs. diffuse cutaneous) and age were utilized in downstream analysis. Gene expression data was probed for senescence signatures as described above. Specifically for aging, Lee et al. found a subset (10 genes) of the established aging genes upregulated in sun exposed skin among publicly available data. (8) These 10 genes were used for the aging analysis in SSc skin.

## 3. Results

Three publicly available gene expression microarray data sets were identified (GSE48149, GSE81292, GSE76808).(29-31) After curation and QC, removing 4 duplicate samples from GSE81292 and 31 non SSc-ILD samples from GSE48149, a total of 38 SSc-ILD subjects and 18 healthy controls were included in the meta-analysis. Studies were successfully merged and batch-corrected (Figure 1B) and the merged gene expression matrix included 11,542 common genes for analysis.

### 3.1 Single gene and pathway analysis reveal markers of cellular senescence and shared features with IPF

There were 586 differentially expressed genes (DEGs) (FC>1.2, q<0.05) between SSc-ILD and healthy controls with 264 genes upregulated, and 322 downregulated in SSc-ILD (Supplementary Table 1). Among the top upregulated DEGs, notable markers included MMP7, a metalloproteinase involved in extracellular matrix (ECM) degradation and biomarker of pulmonary fibrosis, KRT17, an epithelial basal cell marker overexpressed in idiopathic pulmonary fibrosis and SPP1, a biomarker of IPF. GDF15, a marker of cellular stress and senescence, and COMP were also found to be increased (Fig 2A). Hierarchical clustering of the top DEGs revealed separation between SSc-ILD and controls with the emergence of 4 distinct gene clusters. Pathway analysis of individual clusters demonstrating notable down regulation of cell processes, regulation of cell death and angiogenesis while upregulated pathways include ECM formation, epithelial differentiation, collagen organization and senescence (Fig 2B) (Supplementary Table 2). Gene set enrichment analysis revealed enrichment of 8 hallmark pathways including notable enrichment of epithelial mesenchymal transition pathway (FDR q-value < 0.001), coagulation (FDR q-value = 0.02), P53 pathway (FDR q-value = 0.02), and DNA repair (FDR q-value = 0.05) (Figure 2C).

**Figure 2:**
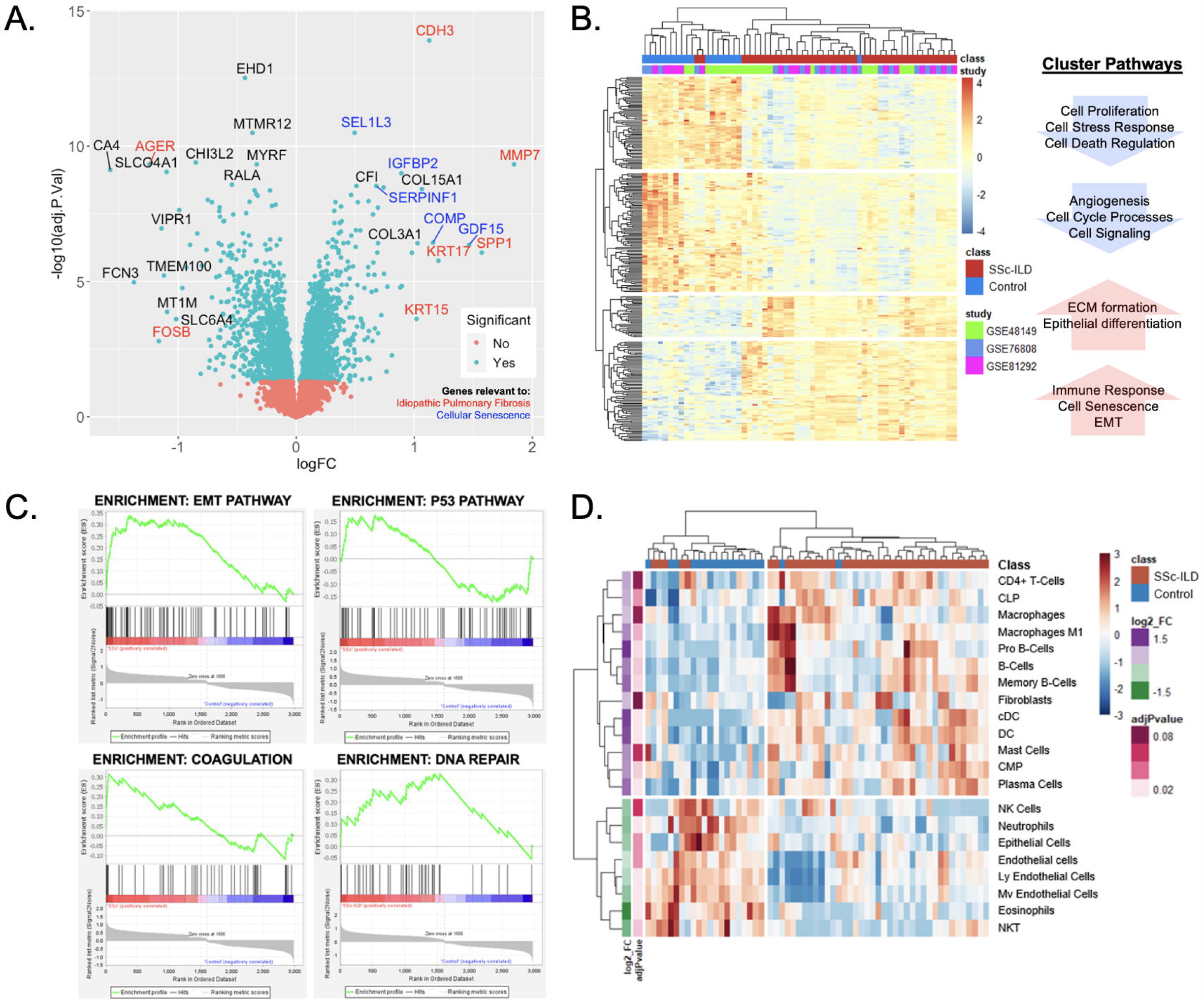
Lung gene expression meta-analysis of SSc-ILD and healthy controls. (A) Volcano plot of significant genes demonstrates overlap with IPF (red) and cellular senescence markers (blue). (B) Heatmap of DEGs with hierarchical clustering demonstrates 4 distinct clusters with associated pathways. (C) Gene set enrichment analysis identifies EMT, p53, coagulation, and DNA repair pathways as most highly enriched in SSc-ILD. (D) Heatmap of cell enrichment analysis revealing increase in dendritic cells and loss of endothelial cells.

### 3.2 Cell enrichment analysis reveals loss of endothelial cells

A total of 45 cell types were examined by cell type enrichment analysis including cells from the epithelial, stromal, and immune cell compartments. 21 cell types were found to be significantly enriched in lung tissue, including 13 enriched in SSc-ILD and 8 enriched in controls (adj p-value < 0.1). Hierarchical clustering demonstrated distinct separation between cases and controls (Figure 2D). B-Cells, dendritic cells, and plasma cells were the most highly increased in SSc-ILD while various types of endothelial cells, neutrophils, eosinophils, and epithelial cells were most significantly depleted in SSc-ILD compared to controls (adj p-value <0.05). Notably fibroblasts were only slightly increased in SSc-ILD.

### 3.3 Aging and senescence pathways are enriched in SSc-ILD

Both cell-specific senescence (DePianto et al) and aging specific (Lee et al.) genes were probed to examine if they were enriched in the SSc-ILD dataset. Among senescence signatures, 145 out of the 227 previously described epithelial genes and 68 out of the previously described 117 fibroblast genes were detected in our dataset. Hierarchical clustering of both epithelial and fibroblast senescence signatures demonstrated a distinct pattern of expression between SSc-ILD patients and healthy controls and found to have a 3.6-fold and 3.7-fold increase respectively in SSc-ILD compared to controls (Figure 3A). When probing the aging gene signature, 8/16 genes upregulated in aging were present in the dataset with 4 significantly increased in SSc-ILD (*CDKN2A, FRZB, PDE1A, NAPI12*) (Figure 3B).

**Figure 3:**
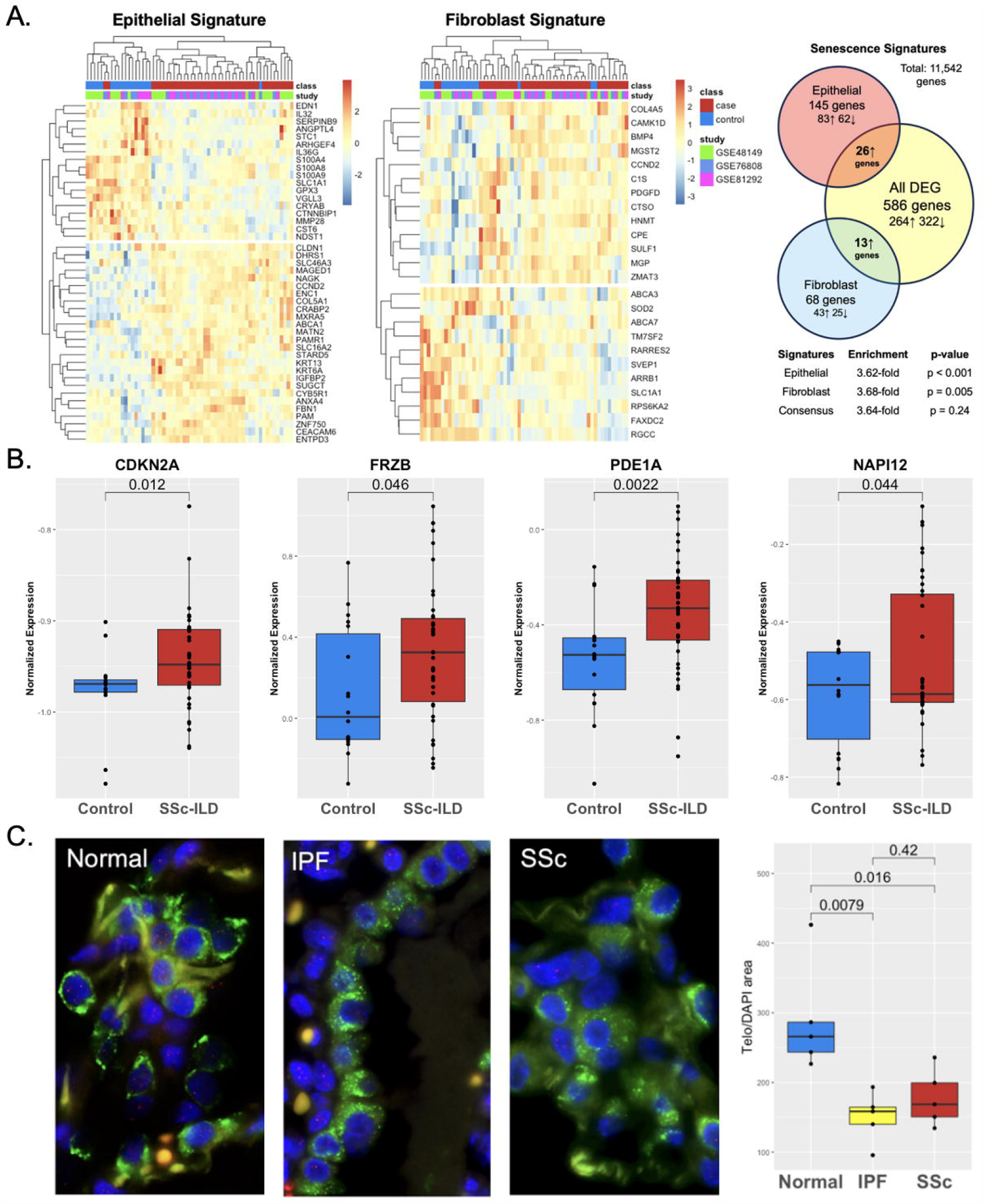
Aging and Senescence are enriched in SSc-ILD. (A) Heatmap of lung epithelial and fibroblast senescence signatures demonstrate distinct patterns of expression in SSc-ILD compared to controls and 3.6-fold, 3.7-fold enrichment respectively. (B) Aging genes are increased in SSc-ILD compared to controls. (C) TeloFISH demonstrates telomere length is shortened in SSc-ILD vs. controls and comparable to IPF.

### 3.4 TeloFISH demonstrates shortened telomere length in SSc-ILD

To validate the aging related findings in the publicly available data, we examined telomere length in lung tissue in SSc-ILD. Given the shared markers and pathways between SSc-ILD and IPF, and lack of understanding of the key pathologic cell type driving SSc-ILD, we focused the telomere studies on type II alveolar cells, the stem cell of the lung epithelium implicated in IPF pathogenesis.(32) Telomere staining was found to be significantly less intense in both SSc-ILD compared age matched control lungs (p = 0.016) and similarly significantly less intense in IPF (p = 0.008). Telomere intensity was similar between IPF and SSc-ILD (p = 0.42) (Figure 3D).

### 3.5 Aging and Senescence signatures are enriched in SSc skin

Both cell-specific senescence and aging specific genes were probed to investigate their presence and relevance in SSc skin including 22 healthy controls and 70 SSc patients (GSE76886). Among senescence signatures, 165 out of 227 epithelial genes and 92 out of 117 fibroblast genes were detected in the skin dataset. Hierarchical clustering of both epithelial and fibroblast senescence signatures demonstrated a distinct pattern of expression between SSc patients and healthy controls, with some separation by SSc cutaneous subtype (limited vs diffuse cutaneous). Notably the senescence signature expression did not cluster by age group (Figure 4A). Compared to all DEGs, the epithelial senescence signature and fibroblast senescence signature was found to have a 2.6-fold and 3.1-fold enrichment respectively in SSc skin disease compared to controls. Among aging specific genes, 10 were reported to be increased in the aging sun exposed skin in publicly available GTEx by Lee et al. Among these genes, 6 were present in our dataset and 4 were significantly increased in SSc skin compared to controls (Figure 4B). Two of the most highly expressed senescence markers in SSc-ILD, GDF15 and COMP, were also significantly increased in SSc skin. Many of the aging and senescent genes demonstrated a stepwise increase between SSc subtypes with diffuse cutaneous having significantly increased expression of *GDF15, CDKN2A* and *PDE1A*, suggesting degree of skin severity may be associated with aging and senescence gene expression.

**Figure 4:**
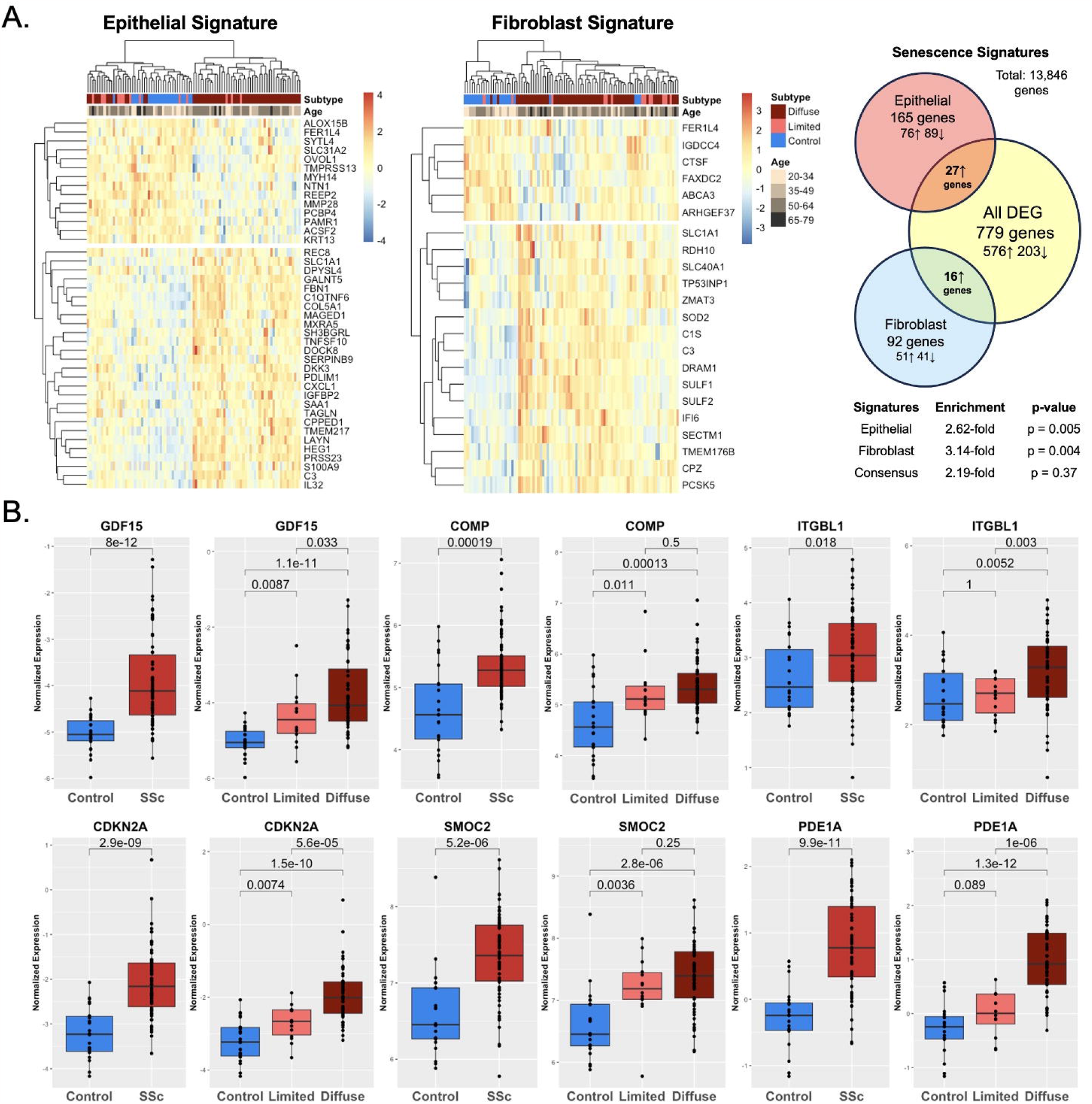
Aging and Senescence are enriched in SSc skin. (A) Heatmap of epithelial and fibroblast senescence signatures demonstrate distinct patterns of expression in SSc skin compared to controls with 2.6-fold, 3.1-fold enrichment respectively. (B) Aging genes are increased in SSc skin compared to controls and expression associated with degree of skin severity (limited vs. diffuse cutaneous).

## 4. Discussion

Advanced age is a known risk factor for fibrotic lung disease and associated with worsened clinical outcomes and decreased survival.(6) Cellular senescence, a hallmark of aging, is a state of replicative arrest leading cells to adopt a senescence-associated secretory phenotype (SASP) which has been shown to have pro-fibrotic effects on the lung.(10) Given their relevance to lung fibrosis pathogenesis, novel aging and senescence signatures have been established and validated in the lung. Our objective was to investigate expression of the signatures in SSc-ILD to ascertain their presence and potential relevance to the disease. Through a gene expression meta-analysis of publicly available data, we demonstrate that markers of senescence (i.e., GDF15, COMP, CDKN2A) and senescence-related pathways (i.e., P53, EMT) are significantly upregulated in SSc-ILD compared to controls. We then took a hypothesis-driven approach and probed established aging and cell-specific senescence signatures and found that both epithelial and fibroblast signatures as well as aging related genes were significantly enriched in lung tissue of SSc-ILD as well as in SSc skin. Finally, the presence of telomere shortening, a hallmark of aging and senescence, in lung epithelium of SSc-ILD further supports the possible mechanistic effects of accelerated aging in SSc-ILD pathogenesis.

Aberrant aging as a driver of pulmonary disease has been well established in several lung diseases including IPF and COPD, however its role in SSc-ILD remains less clear.(11,33,34) Previous studies have reported expression of p53 pathway, a key mediator in apoptosis and senescence, is increased in SSc-ILD as is the presence of p16 in lung tissue of SSc-ILD.(17,19) Telomere length has also been found to be shortened in the peripheral blood of SSc-ILD patients and degree of shortening has been associated with clinical outcomes and survival.(35) In fact, circulating antibodies to telomere associated proteins have been identified in a subset of SSc patients suggesting autoimmunity may contribute to an aberrant aging phenotype.(36) Our findings add to and strengthen the limited but growing body of literature that pathologic aging and senescence may be implicated in SSc-ILD.

Several cell types and mechanisms have been proposed on how cellular senescence may promote the fibrosis of SSc-ILD. For example, endothelial cell senescence has been shown to inhibit angiogenesis and promoted endothelial to mesenchymal transition (EndoMT), which is thought to be profibrotic and can contribute to both dermal and pulmonary fibrosis.(37,38) In our dataset we found there was a significant loss of “normal” endothelial cells by cell enrichment analysis and that angiogenesis pathways were dysregulated in SSc-ILD compared to controls. Similarly, accumulation of senescent fibroblasts, which have increased ECM production and pro-inflammatory cytokine secretion, have been described in SSc, primarily in the skin, as well as murine models of pulmonary fibrosis, and may have a predilection for myofibroblast transformation.(39,40) Additional functional studies are needed to validate aging mediators are drivers of fibrosis in SSc-ILD and importantly, better define whether the fibrosis propagated in the context of aging and cellular senescence stems from the epithelial, stromal, and/or immune compartments.

The role of senescent cells in development of pulmonary fibrosis is well described in IPF and due to the mechanistic and therapeutic implications, there is interest in understanding shared pathology between interstitial lung diseases.(10,17) SSc-ILD has known shared features with IPF including later age of onset, male predilection, and a subset of SSc-ILD patients having identical histopathologic findings of usual interstitial pneumonia (UIP) pattern of fibrosis. In our analysis, we found significant overlapping features between SSc-ILD and IPF, suggesting there may be in part a shared pathobiology between the two diseases. On a single-gene level, MMP7 was the top differentially expressed gene in SSc-ILD. MMP7 is a matrix metalloproteinase responsible for ECM breakdown and well described biomarker in IPF that has shown association with clinical outcomes in IPF.(41) Similarly, SPP1, KRT17 and CDH3, which are upregulated in SSc-ILD, have all been described to be dysregulated in IPF.(42–45) Telomere dysfunction and shortened telomeres are a hallmark of IPF pathogenesis.(11,14,15) We demonstrate telomeres are significantly shortened in SSc-ILD in type II alveolar cells, the pathologic cell type of IPF, compared to controls. ATII cells, which are responsible for maintaining pulmonary homeostasis, when senescent, have been shown to be among the key drivers of the initial fibrosis in IPF. While the roles of ATII cells and other epithelial cells in SSc-ILD need to be elucidated, our data demonstrates telomere attrition is present at the tissue level in SSc-ILD to a similar extent as IPF.

There are limitations to our study. Demographic and clinical characteristic data such as age, sex, medication use, and disease duration were not available on an individual case basis for lung datasets and therefore not utilized for this study. However, each dataset did cite using age similar individuals between cases and controls in their original studies. The sample size is limited and this meta-analysis was cross sectional in nature, providing a snapshot of gene expression in an otherwise dynamic disease. Another limitation is that this study utilized gene expression available from microarray data, given no bulk RNAseq studies present of SSc lung tissue, while the aging and senescence signatures were developed using bulk RNAseq. There were therefore several genes described by these signatures that were not detected in our dataset once merged.

In conclusion, our study contributes to the growing body of evidence that aberrant aging may play a role in SSc-ILD pathogenesis. We demonstrated aging and senescence signatures are upregulated in SSc-ILD and that telomere dysfunction is present in lung tissue of SSc-ILD. These findings have both mechanistic and therapeutic implications, however further studies need to be done. Future work includes examining these signatures on a single cell level to determine the cell type specific transcriptional profiles of senescent cells of interest in SSc-ILD as well as functional studies to understand their implications on disease pathogenesis.

## Funding

This work is in part supported by Rheumatology Research Foundation, NIAMS P30 AR070155, and R01 HL139897.

## Conflict of Interest

*The authors declare that the research was conducted in the absence of any commercial or financial relationships that could be construed as a potential conflict of interest*.

## Notes

### Competing Interest Statement

The authors have declared no competing interest.

